# Direct visualization and characterization of the human zona incerta and surrounding structures

**DOI:** 10.1101/2020.03.25.008318

**Authors:** Jonathan C. Lau, Yiming Xiao, Roy A.M. Haast, Greydon Gilmore, Kamil Uludag, Keith W. MacDougall, Ravi S. Menon, Andrew G. Parrent, Terry M. Peters, Ali R. Khan

## Abstract

The zona incerta (ZI) is a small gray matter region of the deep brain first identified in the 19th century, yet direct *in vivo* visualization and characterization has remained elusive. Noninvasive detection of the ZI and surrounding region could be critical to further our understanding of this widely connected but poorly understood deep brain region and could contribute to the development and optimization of neuromodulatory therapies. We demonstrate that high resolution (submillimetric) longitudinal (T1) relaxometry measurements at high magnetic field strength (7 Tesla) can be used to delineate the ZI from surrounding white matter structures, specifically the fasciculus cerebellothalamicus, fields of Forel (fasciculus lenticularis, fasciculus thalamicus, field H), and medial lemniscus. Using this approach, we successfully derived *in vivo* estimates of the size, shape, location, and tissue characteristics of substructures in the ZI region, confirming observations only previously possible through histological evaluation that this region is not just a space between structures but contains distinct morphological entities that should be considered separately. Our findings pave the way for increasingly detailed *in vivo* study and provide a structural foundation for precise functional and neuromodulatory investigation.

## Introduction

The zona incerta (ZI) is a small but diffuse structure in the deep brain first identified by Auguste Forel in 1877, famously described as “an immensely confusing area about which nothing can be said” (Forel, 1877). Forel appreciated that the ZI consisted of gray matter located between the external medullary lamina of the thalamus and the corpus Luysi (subthalamic nucleus; STN) of otherwise “indefinite” description. It is telling that Forel found the ZI so difficult to describe given his crucial role in the delineation of surrounding white matter tracts still referred to eponymously as the fields of Forel (Gallay et al., 2008). Since its original description, much has been learned about the ZI and surrounding structures although robust *in vivo* visualization has remained elusive.

The anatomical boundaries of the ZI have generally been described in the context of its more discrete neighbors rather than based on any consistent feature of the region itself. Packed in a small area between the ventral thalamus, STN, and red nucleus (RN), the ZI is situated at a complex junction of major white matter pathways including the cerebellothalamic, pallidothalamic, medial lemniscal, and corticospinal tracts. Along its dorsal, ventral, and medial borders, the ZI is surrounded by the fasciculus thalamicus (ft; also known as the H1 field of Forel), the fasciculus lenticularis (fl; also known as the H2 field of Forel), and the H field, which is a convergence of the fl and the ansa lenticularis (al), respectively (Gallay et al., 2008; Nieuwenhuys et al., 2007). The rostral ZI (rZI) is continuous with the reticular nucleus of the thalamus laterally and with the lateral hypothalamus anteromedially. The caudal ZI (cZI) is laterally bounded by the STN and posterior limb of the internal capsule. To date, most of the details regarding the region are the result of meticulous study of post-mortem specimens (Gallay et al., 2008; Morel, 2007; Schaltenbrand and Wahren, 1977).

Cytoarchitectonic and myeloarchitectonic studies in experimental animals (most commonly rodents and primates) have identified the ZI as a gray matter complex consisting of loosely arranged neurons of heterogeneous morphology with a diverse immunohistochemical profile (Nieuwenhuys et al., 2007). In Golgi preparations of the ZI, two main neuronal classes have been identified: principal cells and interneurons (Ma et al., 1997). Gene expression studies have revealed a common embryological origin along with the reticular nucleus of the thalamus and pregeniculate nucleus of the ventral diencephalon, specifically the prethalamic segment, which predominantly contains GABAergic neurons (Puelles et al., 2012; Watson et al., 2014). Through immunohistochemical analysis in experimental animals, a general pattern of at least four component ZI sectors has emerged in the rostral, dorsal, ventral, and caudal directions (Mitrofanis, 2005). Tract-tracing studies have identified extensive and often bilateral connections between the ZI and the cortex, subcortex, and spinal cord (Mitrofanis, 2005; Watson et al., 2014). At least five functional subsectors within the ZI have been suggested: auditory, limbic, motor, somatosensory, and visual. However, unlike other nearby structures like the STN, no robust immunohistochemical biomarker has been described for the ZI proper.

The diversity of chemical expression and widespread connections suggest an important modulatory role of the ZI in regulating brain function. The ZI forms extensive inhibitory connections with spinothalamic relay nuclei in rodents and non-human primates, and thus may play an important role in modulating neuropathic pain and the somatosensory system (Masri et al., 2009; Truini et al., 2013). In a perhaps related manner, in rodent studies, the rostral ZI provides inhibitory control over the thalamus during sleep (Llinás and Jahnsen, 1982; Watson et al., 2014), which may also relate to its perceived role in modulating consciousness (Mitrofanis, 2005; Power and Mitrofanis, 2001). Finally, recent evidence, also in rodents, suggests an important role for the ZI in modulating fear generalization (Venkataraman et al., 2019) and appetite (Zhao et al., 2019).

In humans, the most well-studied role of the ZI is as a putative target for neuromodulatory therapy transmitted either within the cZI or its vicinity, which has been observed to be highly effective for the treatment of essential tremor (Hariz and Blomstedt, 2017). These investigations began in the 1960s with selective ablation (Bertrand et al., 1969; Mundinger, 1965; Spiegel et al., 1962, 1964; Spiegel and Wycis, 1954; Velasco et al., 1975; Wertheimer et al., 1960), but as technologies improved, various groups (Blomstedt et al., 2010; Mohadjer et al., 1990; Nowacki et al., 2018a; Plaha et al., 2006; Velasco et al., 2001) demonstrated that electrical stimulation to these regions was also effective. Yet because of poor direct visualization, controversy has remained as to whether the therapeutic effect is derived from modulation of the cell bodies in the cZI, wayward connections such as the fasciculus cerebellothalamicus (fct; also known as the prelemniscal radiations or raprl) (Castro et al., 2015; Velasco et al., 1972), or some combination of both (Blomstedt et al., 2010). Given the ambiguity and high functional density of the region, the stereotactic target is often considered more broadly as the posterior subthalamic area (PSA) (Blomstedt et al., 2018; Hariz and Blomstedt, 2017; Nowacki et al., 2018a). Targeting of the region relies on identification of the PSA indirectly relative to the adjacent STN and RN, which are visible on T2-weighted (T2w) scans (Blomstedt et al., 2010; Nowacki et al., 2018a).

The increased inherent signal resulting from increasing magnetic field strength has presented an opportunity to visualize brain structures that have not been seen at lower field strengths (DeKraker et al., 2018; Marques and Norris, 2017). Many explorations of the deep brain at 7T have exploited T2w tissue properties enabling visualization of many deep brain nuclei with improved resolution and signal-to-noise ratio (SNR) including the RN, substantia nigra, and STN (Keuken et al., 2013; Plantinga et al., 2018; Schäfer et al., 2012), known to be rich in iron (Haacke et al., 2005; Luigi Zecca et al., 2004). Paralleling these successes, previous attempts at direct visualization of the ZI have focused on the use of T2w contrast, with purported identification of the rZI, but not the cZI (Kerl et al., 2013). In this study, we report that, by employing high-resolution longitudinal (T1) mapping at 7T, robust visualization of the ZI and surrounding WM structures is possible *in vivo* along the entire rostrocaudal axis, allowing comprehensive anatomical characterization of this previously obscure deep brain region.

## Materials and Methods

### Participant and image acquisition details

We recruited 32 healthy participants (46.2 +/- 13.5 years; median: 48 years; range: 20-70 years; 12 female and 20 male; right-handed). This study was approved by the Western University Health Sciences Research Ethics Board (R-17-156). All subjects signed a written consent form to participate. The imaging studies were performed in a 7-Tesla head-only scanner (Siemens Magnetom; Siemens Healthineers, Erlangen, Germany) at the Western University Centre for Functional and Metabolic Mapping (CFMM). An 8-channel parallel transmit/32-receive channel coil was used (Gilbert et al., 2011). After localization and preparatory sequences, each subject underwent a 3D MP2RAGE (Marques et al., 2010), 3D SA2RAGE (Eggenschwiler et al., 2012), and 3D optimized T2w fast-spin echo (T2 SPACE) acquisitions (see Table 1).

**Table 1.** MRI sequence details.

| Sequence |  | TE<br>(ms) | TR<br>(ms) | TI<br>(ms) | Flip<br>Angle (°) | Matrix Size | PAT* | Averages | Resolution<br>(mm<sup>3</sup>) | Acquisition<br>Time (s) |
| --- | --- | --- | --- | --- | --- | --- | --- | --- | --- | --- |
| MP2RAGE | 3D | 2.73 | 6000 | 800/2700 | 4/5 | 342x342x224 | 3 | 1 | 0.7x0.7x0.7 | 10:14 |
| SA2RAGE | 3D | 0.81 | 2400 | 45/1800 | 4/11 | 128x128x64 | 2 | 1 | 1.9x1.9x2.1 | 02:28 |
| SPACE | 3D | 398 | 4000 | NA | variable | 320x320x224 | 3 | 1.6 | 0.7x0.7x0.7 | 10:28 |

\* PAT = parallel acquisition technique (acceleration factor)

### Image pre-processing and template creation

Upon completion of an MRI scan session, the images were pushed to a DICOM server (dcm4che; https://www.dcm4che.org) with automatic data standardization and conversion to the Brain Imaging Data Structure (BIDS) (Gorgolewski et al., 2016) using the autobids platform (https://github.com/khanlab/autobids) deployed on a high-performance compute cluster. Autobids uses scanner-specific heuristics enabled by heudiconv (https://github.com/nipy/heudiconv) preconfigured and validated on multiparametric 7T MRI sequences for DICOM to nifti conversion using dcm2niix (Li et al., 2016) and organization into BIDS.

All individual MRI sequences were corrected for gradient nonlinearities using 3D distortion correction (Glasser et al., 2013; Lau et al., 2018) prior to further processing. The objective of individual preprocessing steps was to adequately prepare the individual MRI sequences for quantitative image analysis and also linear alignment with the subject’s T1-weighted structural MRI scan containerized as BIDS apps (Gorgolewski et al., 2017). The outputs of the preprocessing steps were visually assessed for quality (JL).

### Pre-processing: MP2RAGE

As part of the MP2RAGE acquisition, two different images were created at separate inversions. Using a lookup table, these inversion images were used to create synthetic quantitative T1 maps devoid of proton density contrast, reception field bias, and first order transmit field (B_1_^+^) inhomogeneity. Minimal pre-processing was necessary except for using the B1^+^ field map (SA2RAGE) sequence to correct for intensity inhomogeneity (Eggenschwiler et al., 2012); specifically, no post hoc intensity nonuniformity correction was employed. This SA2RAGE-corrected T1 map was used for quantitative analysis. The T1w image was used as a reference image for rigid-body alignment of the T2SPACE scan.

### Pre-processing: T2SPACE

Raw images from the scanner were observed to have prominent intensity inhomogeneities, which were corrected using an initial nonuniformity correction step with N4 (Sled et al., 1998; Tustison et al., 2010) enabling more accurate registration of the T1w image (and associated brain mask) to T2w. A synthetic T1-T2w fusion image was created by multiplying the T1w by the T2w image (Xiao et al., 2014a) and re-estimating the intensity inhomogeneity again with N4. The original T2w image was denoised using the adaptive non-local means method (Manjón et al., 2010) and the obtained inhomogeneity estimation was applied to the denoised image resulting in a final preprocessed T2w image in the scanner space. Rigid registration to the T1w scan was re-estimated using the preprocessed image. Final preprocessed images included both a T2w volume in the original scanner space as well as one resampled into the T1w structural space. The process was bootstrapped once after creating an initial T2w template (see Section on Template Creation) and using the template for histogram-based intensity normalization. Note that because of the combination of post hoc bias field correction and intensity normalization necessary to produce more homogeneous images, the per voxel values of the T2SPACE images are not directly comparable between scans in a quantitative manner. This processing pipeline has been released and containerized as a BIDS app (https://github.com/khanlab/prepT2space/).

### Template creation

The antsMultivariateTemplateCreation2 pipeline was used for multimodal (T1,T2w) template creation (Avants et al., 2011). A corresponding T2w template (in T1w space) was created after propagating the participant T2w images to T1w template space using the relevant transformations produced using prepT2space. An initial template was created using rigid body alignment of each participant’s T1w scan to the MNI2009bAsym template (0.5 mm isotropic resolution) (Fonov et al., 2009). Over a series of 10 subsequent bootstrapped iterations, the deformable registration (diffeomorphic algorithm) was refined (shrink factors: 12×6×4×2×1; smoothing factors: 6×3×2×1×0vox; max iterations: 100×100×70×50×10; transformation model: Greedy SyN; similarity metric: cross-correlation). Using the derived affine and nonlinear transforms, the individual images (T1 and T2w) were transformed and resampled using trilinear interpolation into the template space. Mean intensity images were generated for each parametric sequence. The log Jacobian was computed, providing an estimate of local deformation required to transform each participant into the template space. The scripts for template creation have been archived for reference. Spatial correspondence was quantified using a recently described anatomical fiducial (AFID) placement protocol with residual AFID registration error (AFRE) being calculated across 32 validated anatomical features (Lau et al., 2019) (RRID:SCR_016623) placed in 3D Slicer (Fedorov et al., 2012) (RRID:SCR_005619). A mapping from our study specific 7T template space to standard MNI coordinates (MNI2009bAsym) has also been provided to facilitate cross-study comparison.

### Region-of-interest segmentation

The ZI, RN, and STN were segmented using the 10th iteration T1 and T2w combined template using ITK-SNAP version 3.6.0. Each rater segmented the regions twice, with sessions spaced more than two weeks apart allowing us to calculate intra- and inter-rater reliability via the Jaccard and Dice coefficients. A representative template segmentation was derived by averaging all segmented ROIs and thresholding by majority voting (>50%) -- this was considered the “gold” standard. Three raters segmented the RN and STN twice using the T2w image (JD, JL, YX). We discovered that substructures of the zona incerta region were also visible, and thus, adopted the nomenclature of Morel (Morel, 2007) for describing the regional anatomy (see Supplementary Table S1 for a glossary of terms and disambiguations). Caudally, the fasciculus cerebellothalamic (fct) and medial lemniscus (ml) could be delineated from the ZI. Rostrally, the fields of Forel, specifically the fasciculus thalamicus (ft or H1 field), fasciculus lenticularis (fl or H2 field), and medially the H field (hf) could also be identified. Each of these structures was segmented twice (two months apart) by the lead author using the T1 template. To our knowledge, these structures have not been previously segmented from *in vivo* images. As such, two stereotactic neurosurgeons (AP, KM) were consulted throughout the ZI segmentation process: first, after the initial segmentations by the lead author (JL); second, after identifying critical boundaries of the ZI particularly rostrally; and finally, to review the final consensus segmentation. Several histological human brain atlases were used as references (Hawrylycz et al., 2012; Mai et al., 2015; Morel, 2007; Schaltenbrand and Wahren, 1977). Consensus segmentations were propagated back into individual subject space using the deformations derived from the template creation step. Accurate spatial correspondence was confirmed by visual inspection by expert raters and also by determining that fiducial registration error was in the millimetric range (Figure 1 and Supplementary Figure S2). Once consensus was achieved, manual segmentations were completed in 5 individual scans and voxel overlap measures using Jaccard and Dice were computed to assess the visibility of individual structures.

**Figure 1.**
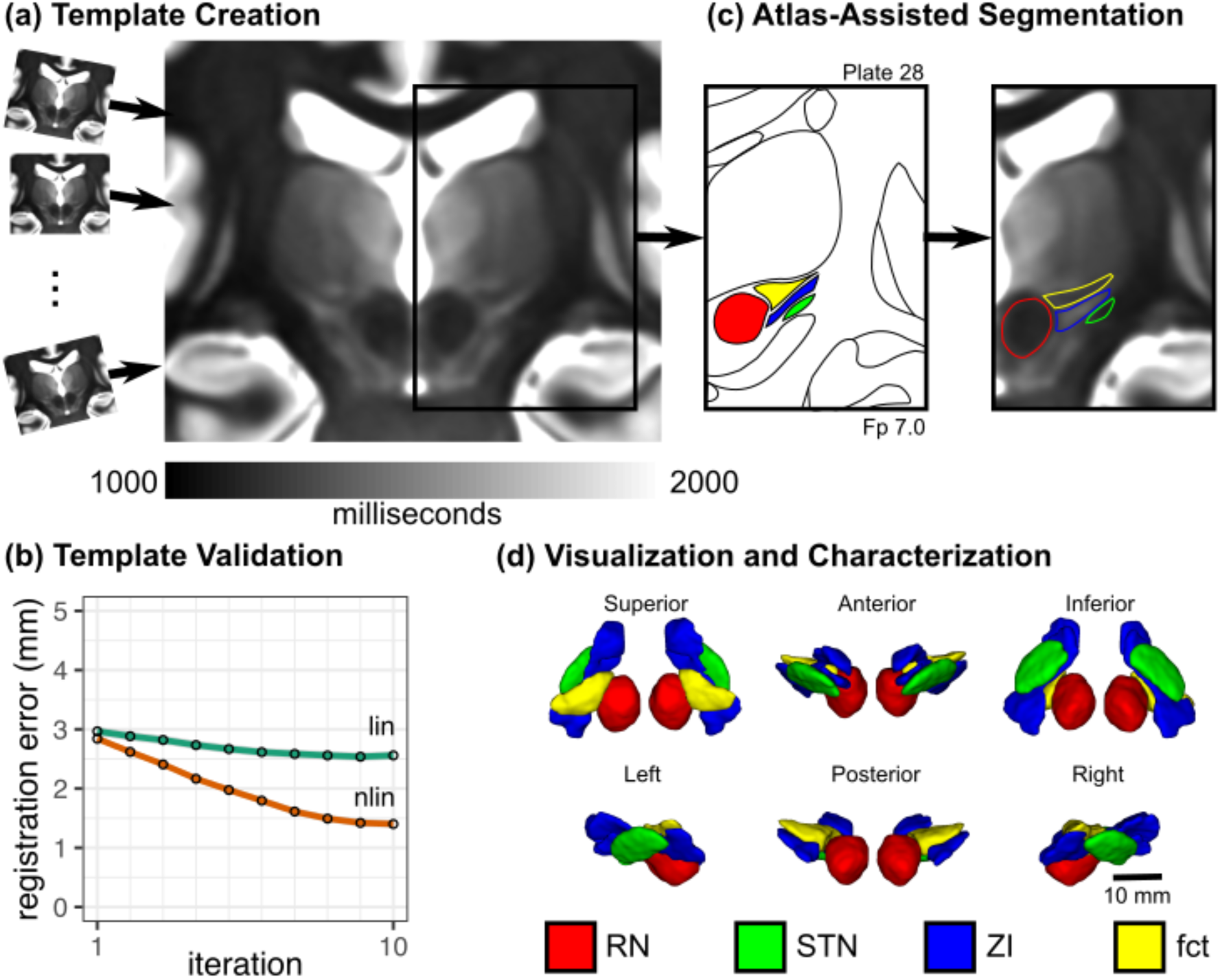
Study workflow for direct visualization and segmentation of the ZI region. (a) To visualize the ZI, we acquired 7T T1 maps from healthy participants. Individual subject data were pooled using deformable template creation methods to create a within-study population average. (b) Registration accuracy stabilized into the millimetric range with increasing complexity of registration (linear to nonlinear) and number of iterations (95% confidence intervals shown; details in Materials and Methods). (c) We found that thresholding our T1 maps to a specific range (1000-2000 ms) revealed similarities with conventional myelin-stained atlases, enabling segmentation of the ZI, demonstrated on Plate 28 (7 mm posterior to MCP) of the Schaltenbrand atlas (Schaltenbrand and Wahren, 1977), corresponding to Plate 48 of the Allen Brain THM Atlas (Hawrylycz et al., 2012) and Plate A6 of the Morel Atlas (Morel et al., 1997). Specifically, the ZI could be distinguished as separate from the fasciculus cerebellothalamicus (fct). Note: Equivalent T1 map images are shown to the left and right of the corresponding Schaltenbrand plate without and with segmentation overlay, respectively. (d) Once consensus segmentations were completed, the structures of the ZI were reconstructed in 3D. Note: the RN and STN labels were segmented based on the corresponding T2w images in this dataset.

The rZI presented some challenges to accurate identification, not for lack of contrast, but due to difficulty with determining its relationship with the fl and ft. On close comparison with histological atlases, the fl appears to run through the rZI. We provide labels for the ZI as a whole, and provide separate labels for the rZI interposed fl, and cZI. Due to partial voluming, the lateral aspect of the central portion of the ZI (between rostral and caudal ends) was too thin to segment along its entire length in our dataset.

### Stereotactic target localization

Three clinicians (two neurosurgeons: KM, AP; one senior neurosurgery resident: JL) placed target locations in the bilateral PSA according to two different placement schemes, which we refer to as Target01 (Blomstedt et al., 2010) and Target02 (Nowacki et al., 2018a) (Figure 4). Both schemes rely on anatomical targeting based on axial T2w images, after performing an initial AC-PC transformation using a validated technique (Lau et al., 2019) in 3D Slicer (Fedorov et al., 2012). Target01 involved the identification of the RN slice of maximal diameter, drawing a horizontal line to mark its equator. The boundary of the STN and its intersection with the RN equatorial line was approximated. Finally, a point was drawn half to two-thirds of the way along the point of STN/RN line intersection and the lateral border of the RN, marking the planned location of the electrode tip. Target02 involved the identification of three different lines: a horizontal line drawn along the equator of the RN identified on the axial slice of maximal diameter, an oblique line drawn along the long-axis of the STN, and finally, an oblique line perpendicular to the long-axis of the STN intersecting the lateral border of the RN at its equator. Consensus placements were agreed upon by the clinicians. The points were placed in the final template space and transformed into the individual participant space. Points in the individual participant space were qualitatively assessed for accuracy.

### Study replication

A second, independent dataset was included to study the inter-site replicability of our findings using age- and sex-matched participant data. This included MP2RAGE and SA2RAGE data acquired at the Maastricht University Brain Imaging Centre (MBIC, Maastricht, Netherlands) using sequence parameters detailed in Table 2 and a 7-T whole-body MRI equipped with a single transmit, 32-channel receive head coil (Nova Medical, Wilmington, MA, USA). Two dielectric pads containing a 25% suspension of barium titanate in deuterated water were placed proximal to the temporal lobe area to locally increase the transmit B1+ field and to improve its homogeneity across the brain (Teeuwisse et al., 2012). Ethical approval for the experimental procedures was provided by the local medical ethics committee (Maastricht University Medical Center, Maastricht, Netherlands). A total of 32 (cognitive) healthy and age-matched participants (46.6 +/- 13.3 years; median: 48.5 years; range: 20-69 years; 17 female and 15 male) were included after obtaining written informed consent.

**Table 2.** MRI sequence details for study replication dataset

| Sequence |  | TE<br>(ms) | TR<br>(ms) | TI<br>(ms) | Flip<br>Angle<br>(°) | Matrix Size | PAT* | Averages | Resolution<br>(mm<sup>3</sup>) | Acquisition<br>Time (s) |
| --- | --- | --- | --- | --- | --- | --- | --- | --- | --- | --- |
| MP2RAGE | 3D | 2.47 | 5000 | 900/2750 | 5/3 | 320x320x240 | 3 | 1 | 0.7x0.7x0.7 | 08:02 |
| SA2RAGE | 3D | 0.78 | 2400 | 45/1800 | 4/10 | 128x128x96 | 2 | 1 | 2x2x2 | 02:16 |

\* PAT = parallel acquisition technique (acceleration factor)

Analysis of the study replication dataset followed a similar workflow as outlined for the primary dataset, including B1+ correction of the MP2RAGE data and template building (see section on Image pre-processing and template creation). In addition, dielectric pads were removed from the images by intensity thresholding the second MP2RAGE inversion image to improve the subsequent template building process. Here, data from two subjects were discarded due to misregistration. Finally, the ZI, rZI, cZI, fct, and the ft segmentations obtained using the primary dataset (see section on Region-of-interest segmentation) were projected onto the replication template by applying a primary-to-replication template registration to allow volumetry and relaxometry analyses in native subject space for evaluation of cross-study use of our segmentations. Manual segmentations were completed in 5 individual scans and voxel overlap measures were computed to assess the visibility of individual structures.

### Direct *in vivo* visualization at standard magnetic fields

Given our findings using high-resolution 7T data, we investigated whether these features could similarly be visible at standard fields, which are more widely accessible. We explored several individual participant datasets using the DESPOT1 (Deoni et al., 2005) and MP2RAGE sequences at standard field. Furthermore, we investigated whether these features were visible on the ICBM MNI2009b template (Fonov et al., 2009) with appropriate windowing, which has been aligned with the BigBrain histological space (Amunts et al., 2013; Xiao et al., 2019).

## Results

The 7T MRI participant data were pooled using deformable template creation methods to create a within-study population average with validation of intersubject spatial correspondence (Lau et al., 2019) (Figure 1; registration accuracy: 1.27 +/- 1.02 mm; Supplementary Figure S2). The population average was reoriented relative to the anterior and posterior commissure allowing coordinates to be expressed relative to the mid-commissural point (MCP). The population averaging technique facilitated further boosting of the contrast and MRI measurements within the ZI region (Figure 1).

### Direct visualization of the human zona incerta and surrounding regions

With appropriate windowing, the contrast from the quantitative T1 maps was highly similar to classic myelin-stained histological atlas (Figure 1) (Schaltenbrand and Wahren, 1977) with white matter structures appearing hypointense relative to surrounding hyperintense gray matter. We found that the human ZI could be directly visualized *in vivo* along its entire rostrocaudal axis as a region of high T1 signal. Moreover, the ZI appeared distinct from the surrounding white matter tracts of the fasciculus thalamicus (ft), fasciculus lenticularis (fl), field H (fh), and medial lemniscus (ml) (Figure 1). Caudally, within the PSA, the fasciculus cerebellothalamicus (fct) could be clearly identified as a distinctly identifiable region of relatively low T1 signal, anterior to the cZI and anterolateral to the RN, a structure previously only identified at high resolution on histological sections (Figures 1c, 4b, 5). Rostrally, regions of high T1 signal were identified both superior and inferior to the fl (Figure 2), leading us to identify an inferior/ventral rZI region ambiguously labeled in existing human atlases (Figure 3b). These substructures were not visible on T2w images.

**Figure 2.**
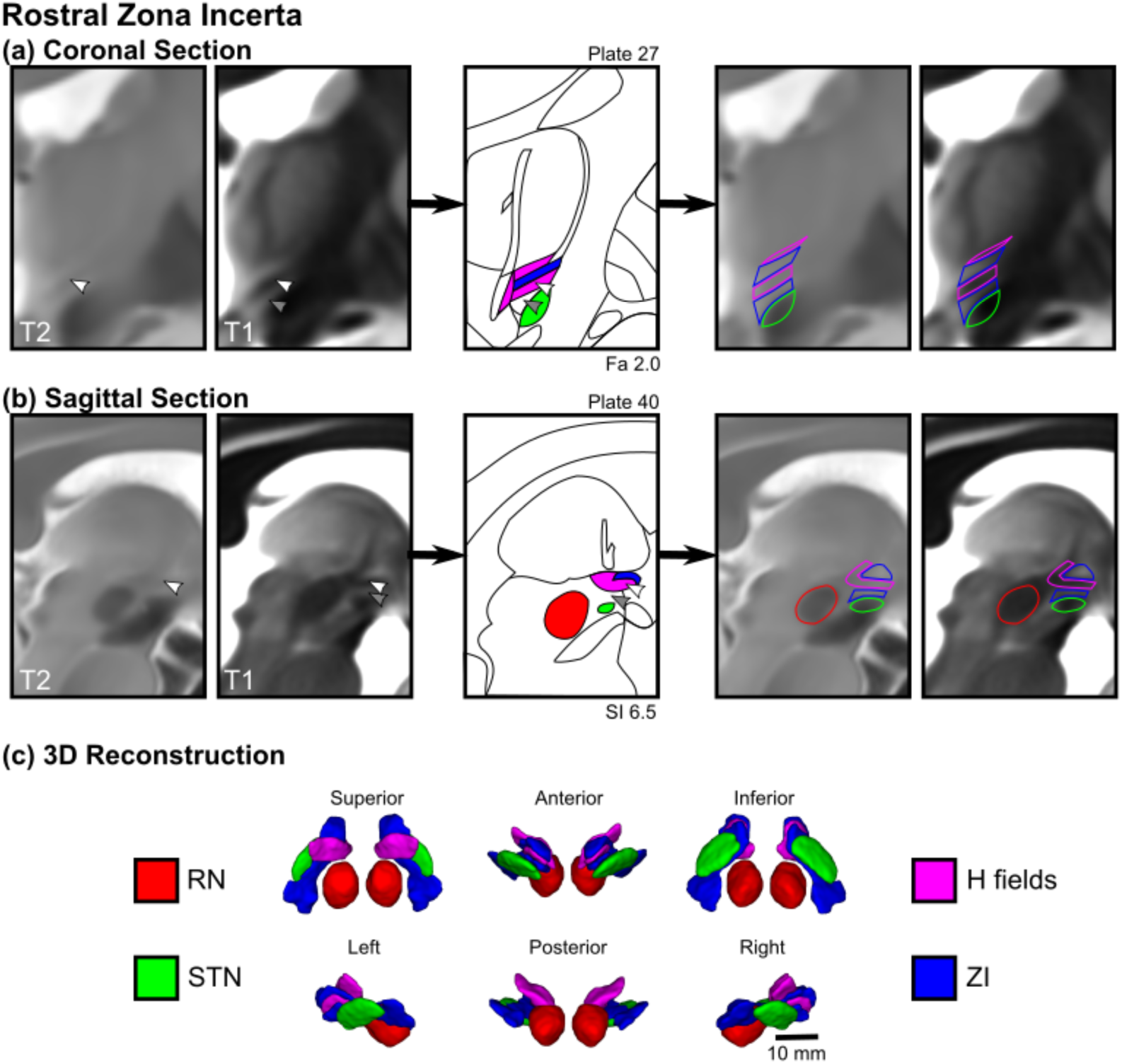
Direct visualization and segmentation of the rostral zona incerta including the fields of Forel. Select views of the rZI demonstrate separate dorsal and ventral components of the rZI as well as the H fields, which include the H1 field (fasciculus thalamicus), H2 field (fasciculus lenticularis), and H field. Equivalent MR images are shown to the left and right of the corresponding Schaltenbrand atlas plate without and with the segmentation overlay, respectively. (a) In the coronal plane, the white arrowhead demonstrates a T2w hypointense region previously identified as the rZI (Kerl et al., 2013). This location is relatively T1 hypointense, corresponding spatially and in terms of tissue characteristics to the myelinated H2 field (fl). Below this region (gray arrowhead) is an unlabeled T1 hyperintense region of the Schaltenbrand atlas (Plate 27; 2.0 mm anterior to MCP), corresponding to Plate 39 of the Allen Brain THM Atlas (Hawrylycz et al., 2012) and Plate A13 of the Morel Atlas (Morel et al., 1997). This location corresponds with the ventral rZI identified in other species (Mitrofanis, 2005). (b) These features are similarly identified in the sagittal view with a corresponding representative histological slice from Schaltenbrand (Plate 40. 6.5 mm lateral to MCP), corresponding to Plate L9.1 (6.3) of the Morel Atlas. Note: the Schaltenbrand atlas represents separate post-mortem specimens in each cardinal orientation (coronal, sagittal, axial).

**Figure 3.**
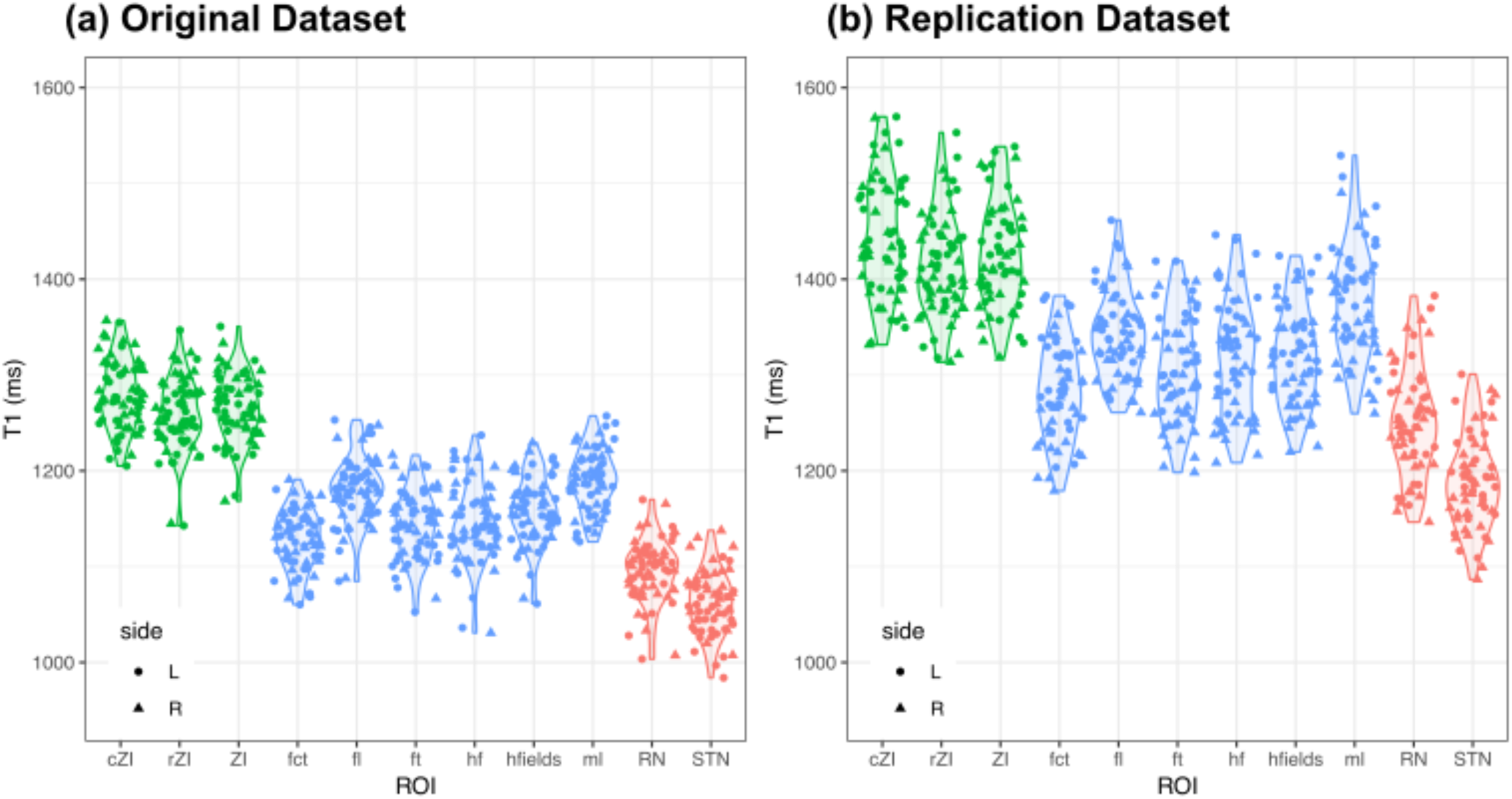
Distinct T1 values for different substructures of the zona incerta region as determined in our original dataset and a replication dataset. The general trends are the same with statistically significant differences in T1 values between the ZI (green) and surrounding white matter (blue) and gray matter (red) regions. The differences between datasets is an observed phenomenon from other studies related to inter-scanner differences and reviewed in the Discussion and a recent study (Haast et al., 2020). Although different, our analysis demonstrates that for a given scanner these tissue characteristics are relatively precise and allow the separation of these regional structures on the basis of local MRI characteristics alone.

**Figure 4.**
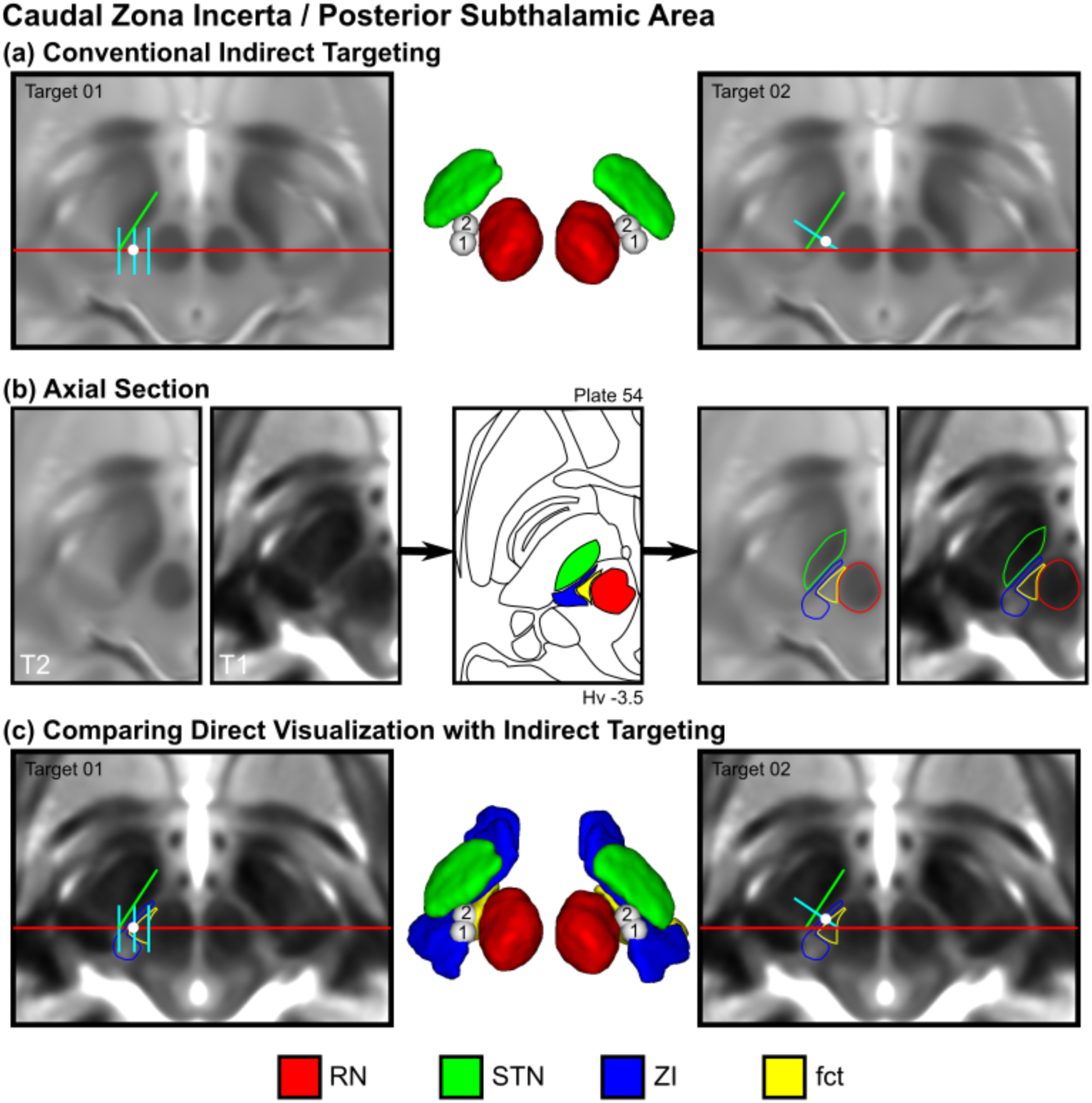
Direct visualization of conventional indirect targets of the posterior subthalamic area. (a) Two conventional indirect targeting methods, Target01 (Blomstedt et al., 2010) and Target02 (Nowacki et al., 2018a), used for stereotactic targeting in the PSA based on the relative location of the RN and STN at the level of the maximal diameter of the RN. (b) Using the Schaltenbrand atlas as a reference (Plate 54, 3.5 mm below MCP), also corresponding to Plate V2.7 of the Morel atlas, we identified the cZI and fasciculus cerebellothalamicus (fct) using the T1 maps (thresholded between 1000-2000 ms). Equivalent MR images are shown to the left and right of the corresponding Schaltenbrand atlas plate without and with the segmentation overlay, respectively. (c) Using the T1 map as an underlay image for our indirect targets provides additional detail regarding the location of the target relative to the cZI and fct, demonstrating that the targets are at the boundary between the two structures. Note: the 3D reconstructions represent an inferior surface view of the ZI region, which best depicts the location of the targets relative the surrounding structures.

**Figure 5.**
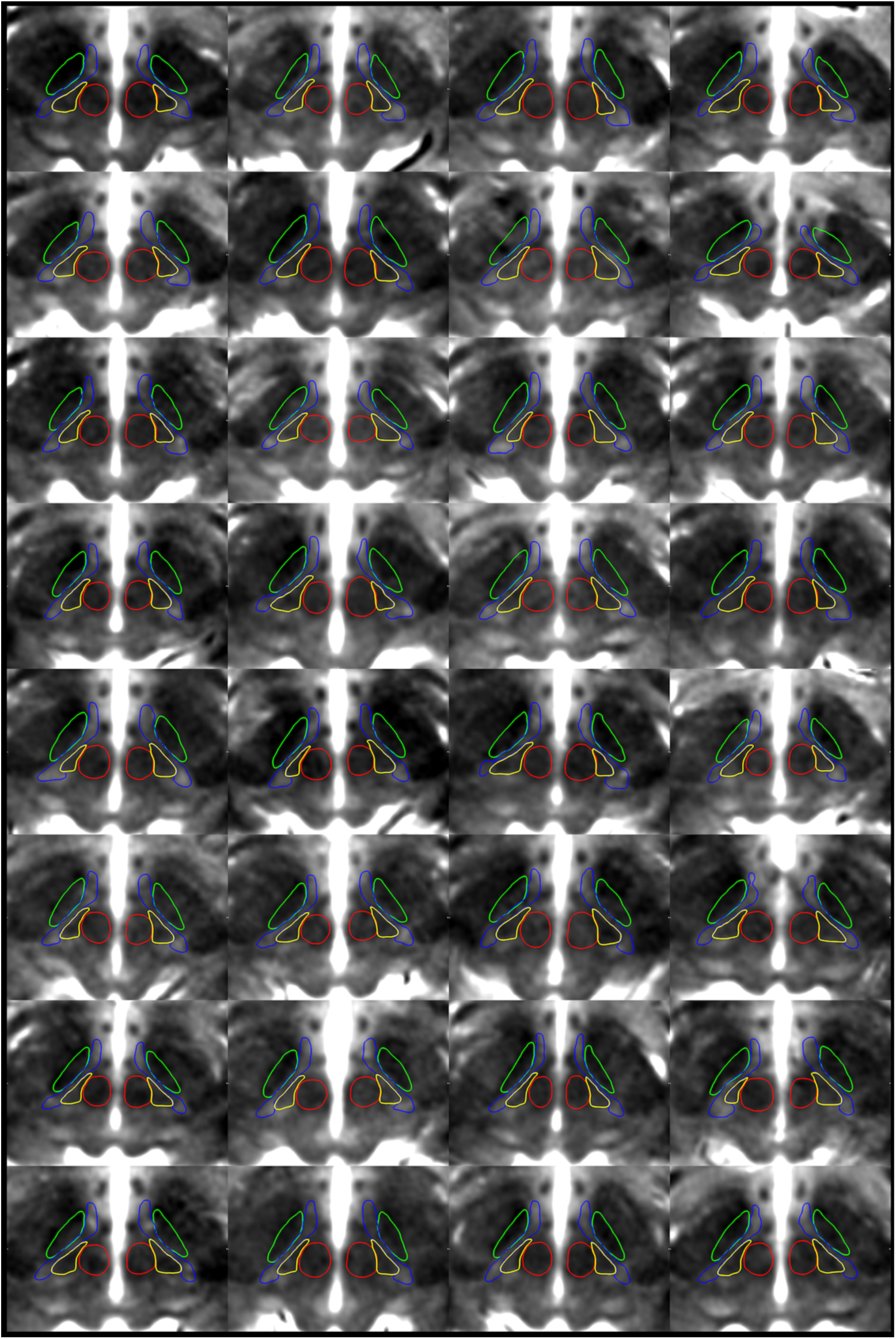
Montage of participant data demonstrating the ability to delineate the zona incerta substructures using high-resolution T1 maps. Representative axial slices demonstrate the ability to distinguish between the fct (yellow) and ZI (blue) at the level of the RN (red) for each individual participant (N=32). Note the RN and STN (green) were initially segmented using the T2w images for the same subject fused into T1 space. The T1 maps are thresholded between 1000-2000 ms.

### Characterization of the human zona incerta and surrounding regions

Direct visualization afforded us an opportunity to segment and characterize the human ZI morphologically using methods not previously possible (see Materials and Methods for details). The segmentations could generally be performed reliably (Dice scores > 0.7) in both the template space and for individual datasets, with details provided in Table 3 and S3. As expected, manual segmentations could be performed more reliably in the template space. For individual subjects, the ZI, cZI, and rZI were segmented in individual subjects with a mean Dice score of 0.72, 0.76, and 0.68, respectively. The surrounding white matter structures (fct, fl, ft, hf, hfields, and ml) were segmented in individual subjects with a mean Dice score of 0.71, 0.63, 0.69, 0.71, 0.71, and 0.73, respectively. The STN and RN mean Dice scores were 0.91 and 0.78, respectively, consistent with previous studies.

**Table 3.** Summary of voxel overlap measures for manual segmentations of the original and replication dataset.

|  |  | Template Segmentation |  | Individual Subject Segmentation |  |  |  |
| --- | --- | --- | --- | --- | --- | --- | --- |
|  |  | Original Dataset |  | Original Dataset |  | Replication Dataset |  |
| Region | Side | jaccard | kappa | jaccard | kappa | jaccard | kappa |
| cZI | L | 0.88±0.01 | 0.94±0.01 | 0.63±0.04 | 0.77±0.03 | 0.64±0.04 | 0.78±0.03 |
|  | R | 0.84±0.02 | 0.91±0.01 | 0.59±0.02 | 0.74±0.02 | 0.59±0.04 | 0.74±0.03 |
|  | combined | 0.86±0.03 | 0.93±0.02 | 0.61±0.03 | 0.76±0.03 | 0.62±0.05 | 0.76±0.04 |
| fct | L | 0.80±0.14 | 0.89±0.09 | 0.55±0.01 | 0.71±0.01 | 0.53±0.03 | 0.69±0.03 |
|  | R | 0.80±0.14 | 0.88±0.08 | 0.55±0.01 | 0.71±0.01 | 0.56±0.06 | 0.72±0.05 |
|  | combined | 0.80±0.11 | 0.89±0.07 | 0.55±0.01 | 0.71±0.01 | 0.55±0.05 | 0.71±0.04 |
| fl | L | 0.76±0.15 | 0.86±0.09 | 0.45±0.02 | 0.62±0.02 | 0.46±0.07 | 0.63±0.07 |
|  | R | 0.79±0.14 | 0.88±0.09 | 0.47±0.02 | 0.64±0.01 | 0.45±0.07 | 0.61±0.07 |
|  | combined | 0.77±0.12 | 0.87±0.08 | 0.46±0.02 | 0.63±0.02 | 0.45±0.07 | 0.62±0.07 |
| ft | L | 0.79±0.01 | 0.89±0.01 | 0.53±0.03 | 0.69±0.02 | 0.47±0.07 | 0.63±0.07 |
|  | R | 0.80±0.03 | 0.89±0.02 | 0.52±0.02 | 0.68±0.02 | 0.48±0.08 | 0.64±0.07 |
|  | combined | 0.80±0.02 | 0.89±0.01 | 0.53±0.02 | 0.69±0.02 | 0.47±0.07 | 0.64±0.07 |
| hf | L | 0.79±0.06 | 0.88±0.04 | 0.55±0.04 | 0.71±0.04 | 0.53±0.07 | 0.69±0.06 |
|  | R | 0.79±0.07 | 0.88±0.04 | 0.56±0.01 | 0.72±0.01 | 0.55±0.05 | 0.71±0.04 |
|  | combined | 0.79±0.06 | 0.88±0.03 | 0.55±0.03 | 0.71±0.03 | 0.54±0.06 | 0.70±0.05 |
| hfields | L | 0.81±0.04 | 0.89±0.02 | 0.55±0.02 | 0.71±0.02 | 0.53±0.05 | 0.69±0.04 |
|  | R | 0.82±0.03 | 0.90±0.02 | 0.56±0.02 | 0.72±0.01 | 0.54±0.04 | 0.70±0.04 |
|  | combined | 0.81±0.03 | 0.90±0.02 | 0.56±0.02 | 0.71±0.02 | 0.53±0.04 | 0.70±0.04 |
| ml | L | 0.82±0.20 | 0.89±0.12 | 0.57±0.02 | 0.72±0.02 | 0.52±0.05 | 0.68±0.04 |
|  | R | 0.86±0.01 | 0.92±0.01 | 0.58±0.02 | 0.74±0.02 | 0.53±0.08 | 0.69±0.07 |
|  | combined | 0.84±0.12 | 0.91±0.07 | 0.58±0.02 | 0.73±0.02 | 0.53±0.06 | 0.69±0.05 |
| rZI | L | 0.78±0.14 | 0.87±0.09 | 0.51±0.02 | 0.68±0.01 | 0.52±0.04 | 0.68±0.03 |
|  | R | 0.79±0.11 | 0.88±0.07 | 0.51±0.01 | 0.67±0.01 | 0.51±0.05 | 0.68±0.04 |
|  | combined | 0.78±0.11 | 0.88±0.07 | 0.51±0.01 | 0.68±0.01 | 0.52±0.04 | 0.68±0.04 |
| ZI | L | 0.83±0.09 | 0.91±0.05 | 0.56±0.01 | 0.72±0.01 | 0.57±0.03 | 0.73±0.03 |
|  | R | 0.83±0.08 | 0.90±0.05 | 0.55±0.01 | 0.71±0.01 | 0.55±0.05 | 0.71±0.04 |
|  | combined | 0.83±0.07 | 0.91±0.04 | 0.56±0.01 | 0.72±0.01 | 0.56±0.04 | 0.72±0.04 |
| RN* | L | 0.95±0.03 | 0.98±0.02 | 0.82±0.05 | 0.90±0.03 |  |  |
|  | R | 0.95±0.03 | 0.98±0.01 | 0.84±0.02 | 0.91±0.01 |  |  |
|  | combined | 0.95±0.03 | 0.98±0.01 | 0.83±0.04 | 0.91±0.02 |  |  |
| STN* | L | 0.90±0.10 | 0.94±0.06 | 0.64±0.06 | 0.78±0.04 |  |  |
|  | R | 0.89±0.10 | 0.94±0.05 | 0.65±0.03 | 0.78±0.02 |  |  |
|  | combined | 0.89±0.10 | 0.94±0.05 | 0.64±0.04 | 0.78±0.03 |  |  |

\* The RN and STN were segmented using the T2w scan, which was not acquired in the replication dataset.

Three-dimensional reconstructions permitted the identification of the ZI as an elongated band situated along the long axis of the STN with broader and more prominent components extending both rostrally and caudally (Figure 1d). To provide a sense of scale, the total volume described here represents a region with a bounding box of 20 mm x 10 mm x 10 mm or smaller than the tip of an adult human finger. Our analysis permitted the identification of concrete dimensions of the ZI, which spans on average approximately 20.4 mm along its main axis (rostrocaudally), 7.4 mm maximally along its secondary axis (medial to lateral), and varying in thickness from less than 1.0 mm along its lateral boundary to 3.6 mm in the cZI (Figure 1d). Calculations of rostral thickness were complicated by the wayward fasciculus lenticularis (see previous section; Figure 2), which if included, is as thick as 7.0 mm, whereas the dorsal rZI and ventral rZI have thickness of 3.7 mm and 1.8 mm, respectively when considered separately. The volume of the ZI was 252.4±22.4 mm^3^ with caudal and rostral components 83.6±8.7 and 169.2±16.3 mm^3^ respectively.

Morphological characterization could be extended to surrounding gray and white matter regions given they could also be well visualized. The RN and STN have been well-characterized in previous studies (Keuken et al., 2017; Xiao et al., 2014b), providing anatomical boundaries to the ZI region with reliability consistent with prior studies (Supplementary Table S3a and S3b). Volumetric results were consistent with previous studies of the RN (296.4±27.8 mm^3^) and the STN (138.9±14.0 mm^3^). Of particular note, the fct was 12.2 mm along its longest axis, 5.4 mm (medial to lateral), and 5.0 mm thick maximally (medially) with a total volume of 135.7±13.3 mm^3^, similar in volume to the STN (138.9±14.0 mm^3^), but located more posteriorly, and separated by the interposed gray matter of the middle to caudal ZI. The fields of Forel (fasciculus thalamicus, fasciculus lenticularis, field H) similarly could be distinguished from the rZI and separately segmented based on differences in T1 intensity with a total volume of 153.7±15.9 mm^3^, also with a volume similar to the STN. The fl and ft tracts themselves formed concentrated bundles of around 1.2 mm diameter and could be distinguished anatomically as separate from the ZI with total volumes 52.0±5.7 mm^3^ and 84.3±8.8, respectively, with the H field medially, consisting of the mergence of the ft and al tracts (Gallay et al., 2008), being 54.6±5.7 mm^3^. The locations of the structures in reference to the MCP (Table 4) were consistent with known values.

**Table 4.** Summary of volume, T1 values, and location relative to the MCP for the zona incerta and surrounding structures.

|  |  |  |  | Coordinates (mm) |  |  |
| --- | --- | --- | --- | --- | --- | --- |
| Region | Side | Volume (mm<sup>3</sup>) | T1 (ms) | x | y | z |
| cZI | L | 87.4±8.1 | 1272.9±34.7 | -12.42±0.78 | -8.04±1.15 | -5.16±1.18 |
|  | R | 79.8±7.6 | 1286.6±36.5 | 12.59±0.81 | -7.68±1.21 | -5.12±1.09 |
|  | *combined* | 83.6±8.7 | 1279.7±36.0 |  |  |  |
| rZI | L | 168.7±16.6 | 1250.4±36.8 | -6.97±0.54 | 1.69±0.53 | -0.79±0.53 |
|  | R | 169.8±16.3 | 1266.0±36.9 | 7.11±0.61 | 2.44±0.61 | -0.88±0.59 |
|  | *combined* | 169.2±16.3 | 1258.2±37.4 |  |  |  |
| ZI<sup>a</sup> | L | 254.3±22.5 | 1258.6±34.4 | -8.82±0.56 | -1.61±0.63 | -2.27±0.52 |
|  | R | 250.6±22.5 | 1272.3±34.5 | 8.86±0.61 | -0.81±0.73 | -2.28±0.46 |
|  | *combined* | 252.4±22.4 | 1265.4±34.8 |  |  |  |
| fct | L | 136.0±12.8 | 1120.5±30.5 | -10.24±0.68 | -5.24±0.78 | -2.11±0.83 |
|  | R | 135.4±14.0 | 1135.3±29.1 | 10.00±0.67 | -4.83±0.88 | -2.62±0.73 |
|  | *combined* | 135.7±13.3 | 1127.9±30.5 |  |  |  |
| fl | L | 51.8±5.9 | 1178.9±34.9 | -6.09±0.56 | 3.36±0.50 | -0.49±0.65 |
|  | R | 52.2±5.6 | 1181.4±34.5 | 6.65±0.64 | 3.81±0.63 | -0.51±0.76 |
|  | *combined* | 52.0±5.7 | 1180.2±34.4 |  |  |  |
| ft | L | 84.0±9.2 | 1135.2±35.6 | -7.24±0.59 | -0.12±0.60 | 1.31±0.43 |
|  | R | 84.5±8.6 | 1150.9±33.4 | 7.58±0.69 | 0.39±0.73 | 1.14±0.46 |
|  | *combined* | 84.3±8.8 | 1143.1±35.1 |  |  |  |
| hf | L | 53.4±5.5 | 1142.8±41.8 | -5.00±0.47 | -0.08±0.52 | -1.24±0.38 |
|  | R | 55.7±5.6 | 1148.2±40.9 | 5.13±0.53 | 0.39±0.62 | -1.28±0.42 |
|  | *combined* | 54.6±5.7 | 1145.5±41.1 |  |  |  |
| hfields<sup>b</sup> | L | 153.4±16.2 | 1154.1±35.0 | -6.46±0.55 | 1.13±0.53 | 0.29±0.46 |
|  | R | 154.0±15.8 | 1162.2±34.1 | 6.83±0.63 | 1.55±0.65 | 0.16±0.51 |
|  | *combined* | 153.7±15.9 | 1158.1±34.5 |  |  |  |
| ml | L | 38.6±3.8 | 1188.4±34.6 | -9.04±0.73 | -10.58±0.91 | -3.39±1.42 |
|  | R | 29.2±3.0 | 1192.2±27.7 | 8.96±0.77 | -10.14±1.03 | -3.40±1.31 |
|  | *combined* | 33.9±5.8 | 1190.3±31.2 |  |  |  |
| RN | L | 292.1±27.1 | 1093.2±33.4 | -4.56±0.48 | -5.92±1.05 | -6.56±0.89 |
|  | R | 300.7±28.3 | 1096.6±32.4 | 4.53±0.52 | -5.65±1.09 | -6.54±0.84 |
|  | *combined* | 296.4±27.8 | 1094.9±32.7 |  |  |  |
| STN | L | 144.2±13.3 | 1045.8±27.3 | -10.00±0.66 | -0.47±0.79 | -3.15±0.57 |
|  | R | 133.6±12.7 | 1078.2±29.2 | 10.23±0.70 | -0.01±0.88 | -3.29±0.57 |
|  | *combined* | 138.9±14.0 | 1062.0±32.4 |  |  |  |

<sup>a</sup> includes rZI and cZI
<sup>b</sup> includes fl, ft, hf

T1 measurements facilitated the identification of substructures of the ZI region in ways that T2w images did not (Table 4). Notably, in the main reported dataset, T1 values were robustly in the 1200-1300 ms range (1265.4±34.8 ms) in the ZI (Table 4 and Figure 3). No differences in T1 values were found between the rZI and cZI (p-value n.s.). To examine whether peri-zonal substructures could be distinctly separated as suggested from our qualitative observations, we compared the T1 values in the ZI against the values in surrounding regions. Wilcoxon rank testing confirmed that T1 mapping was effective at distinguishing the ZI from surrounding local structures (Figure 3). The RN and STN had demonstrably shorter T1 times than the ZI (1094.9±32.7 and 1062.0±32.4 ms respectively; p-value < 0.01). The surrounding white matter tracts were also clearly separable from the ZI, despite their small size, due to distinctly shorter T1 times. Such tracts include the fields of Forel, the fasciculus lenticularis (fl) inferiorly (volume: 52.0±5.7 mm^3^; T1: 1180.2±34.4 ms, p < 0.01) and the fascicularis thalamicus (ft) superiorly (volume: 84.3±8.8 mm^3^; T1: 1143.1±35.1 ms, p < 0.01). The fasciculus cerebellothalamicus (fct) was also distinct from the ZI (volume: 252.4±22.4 mm^3^; T1: 1127.9±30.5 ms), which is of relevance to known surgical neuromodulatory targets.

These analyses were repeated in a matched dataset from Maastricht University (see Materials and Methods; collaborators: RH and KU). Wilcoxon rank sum testing again confirmed intensity differences between the ZI and neighboring white matter and gray matter structures (p-value < 0.01). These results are reported in detail in the associated notebook provided on GitHub. Although the exact T1 values differed (Figure 3), intra-regional variability in T1 was comparably low across datasets (Figure 3b). These inter-scanner differences are a known phenomenon explored in a recent study (Haast et al., 2020).

### Direct evaluation of indirect surgical targets of the zona incerta region

Surgical targets of the ZI region have conventionally been targeted indirectly and the specific area that results in a therapeutic effect remains controversial. We used the high-resolution combined T1 and T2w *in vivo* maps reported in this study to directly evaluate two conventional indirect targets located in the ZI region. Specifically, the PSA is targeted using features from T2w contrast based on the relative positions of the STN and RN. We have demonstrated that the cZI and fct can be separated on the basis of both anatomical location and underlying T1-based tissue characteristics. Two commonly described indirect targets, here referred to as Target01 (Blomstedt et al., 2010) and Target02 (Nowacki et al., 2018a), were placed on T2w images, allowing us to evaluate to which feature (or features) this best corresponded on our T1 maps (Figure 4).

The target placements anatomically corresponded to the boundary between the cZI and the fct lateral to the ipsilateral RN (Figure 4c). This observation was quantitatively supported by our finding that mean T1 values at the surgical targets were lower than in the cZI for both Target01 and Target02 (Supplementary Figure S4c; Wilcoxon rank sum testing p < 0.05), but higher than values in the fct. We also calculated the distance between each indirectly placed target and separately the centroid of the ipsilateral fct and cZI, and assessed whether the indirect target was closer to one or the other. Target01 was almost equidistant from the cZI and fct (3.30±0.22 vs 3.46±0.27 mm respectively), while Target02 was further from cZI and closer to fct (4.62±0.36 vs 2.64±0.52 mm respectively). Differences were confirmed using Wilcoxon rank testing (p < 0.01; Supplementary Figure S4a and S4b). The ability to separate the fct from the cZI is demonstrated pictorially in a montage of all participant data (Figure 5) as well as quantitatively (Figure 3).

### The zona incerta region at standard magnetic field strength

Motivated by our discovery at 7T, we investigated whether we could identify a similar feature at standard magnetic fields, given that they are more widely accessible. Indeed, we determined that on individual T1 map datasets at 3T, a region of relative hypointensity could be seen that represents the gray matter regions of the ZI (Figure 6a). Thus, for practical purposes, a properly optimized T1 map protocol may be sufficient for identification of the nuclear region. In addition, we determined that the ZI is visible on T1-weighted images as a relatively hypointense feature (Figure 6b), when windowed, although the windowing values themselves are arbitrary. Finally, we transformed our regions into the MNI2009bAsym space for use by other groups, which has the advantage of also having close correspondence with the BigBrain template (Xiao et al., 2019).

**Figure 6.**
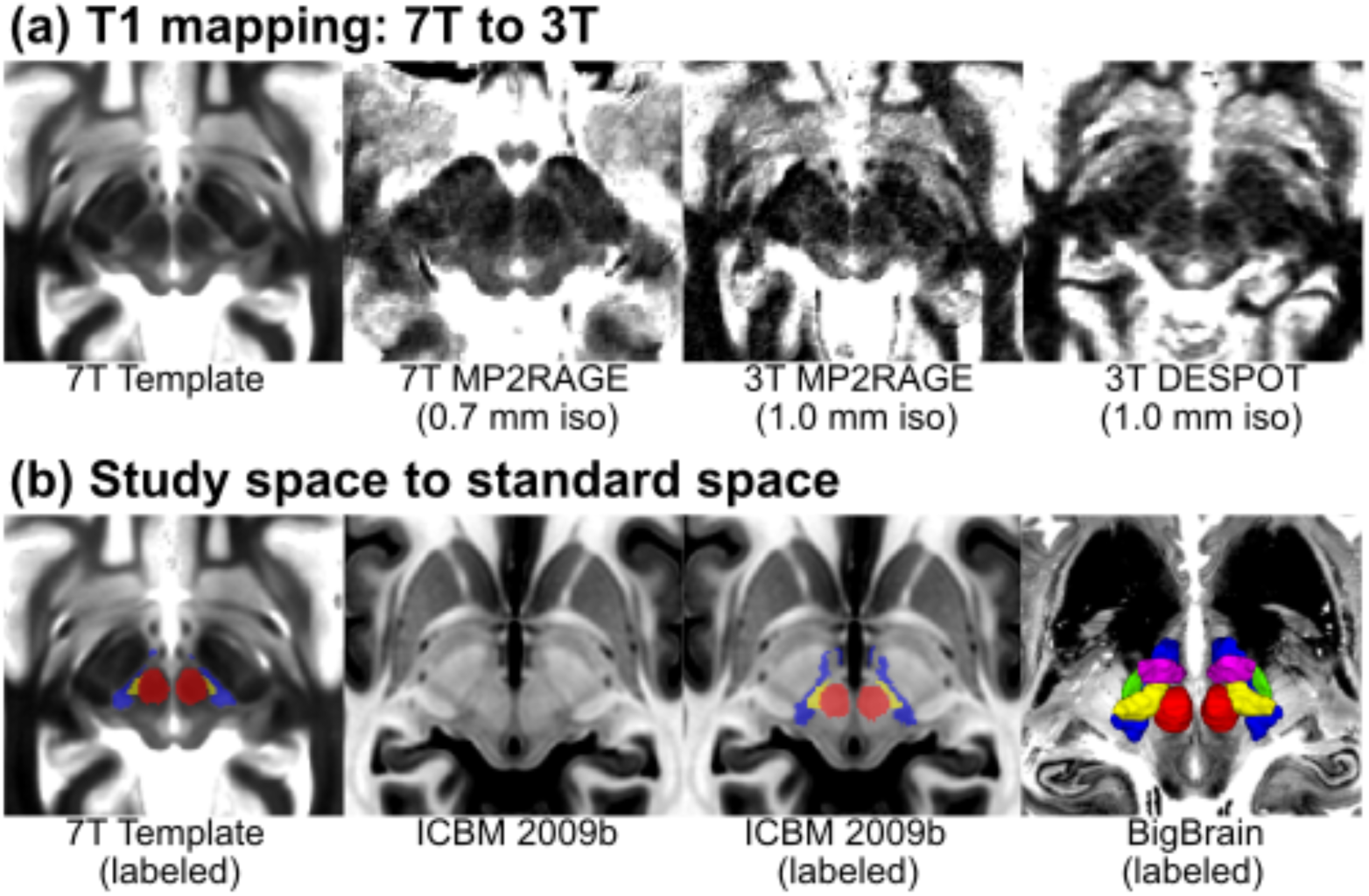
Implications for standard magnetic field strengths and standard spaces. (a) Based on our findings, we investigated whether the ZI substructures could be visualized at standard magnetic field strength. Based on our qualitative assessment, the cZI could be well visualized even at 3T using two different techniques (1.0 mm isotropic resolution, compared to 0.7 mm isotropic resolution at 7T). (b) The ZI subregions created in this study have been transformed into ICBM 2009b space to benefit the neuroscience community. Note that on the T1w ICBM 2009b template, the cZI appears as a hypointense region while the fasciculus cerebellothalamicus is relatively hyperintense (inverse of T1 map). However, the units in the ICBM space are arbitrary. The 3D reconstructions are overlaid on top of the BigBrain template (Amunts et al., 2013; Xiao et al., 2019).

### Data Availability Statement

The template data have been deposited on the Open Science Framework website (https://osf.io/c8p5n/). Code to reproduce this analysis is available at https://github.com/jclauneuro/zona-analysis/. Videos in the study template space are provided with the main labels in each of the standard orientations as Supplementary Material S6.

## Discussion

The present study demonstrates that robust visualization of the ZI and surrounding structures is possible using high-resolution quantitative T1 mapping. We report the first precise delineation of the ZI region *in vivo* providing estimates of the morphology (volume, dimensions) and T1 values. We found that the T1 relaxometry parameters of the ZI were distinct from surrounding white matter pathways. This finding enabled us to decouple a component of the rZI as separate and inferior to the fascicularis lenticularis, which to our knowledge has not previously been labeled on histological atlases of the human brain (Figure 2). Due to the striking similarity in tissue contrast with classic post-mortem myelin staining, we were able to segment the fct as a substructure within the PSA separate from the cZI. This methodology was then used for prospective identification of the active stimulation location for deep brain stimulation for which current standard-of-care relies on indirect targeting.

Efforts at visualizing small structures of the deep brain using high-field MRI have mostly focused on T2w relaxation properties due to the high paramagnetic contrast produced by many subcortical nuclei due to endogenous ferritin (Haacke et al., 2005; Rudko et al., 2014; L. Zecca et al., 2004). Increasing the strength of the main magnetic field (B_0_) results in an at least linear increase in signal, a two- to three-fold increase compared to conventional clinical field strengths. This increased signal can be exploited in a number of ways, including higher resolution (submillimetric) imaging. Visualization at high fields has led to more robust imaging of small structures including the STN and SN using T2w contrast mechanisms (Keuken et al., 2013). The ZI has proven to be elusive to visualization using T2w contrast. In one study at 7T, using a T2w sequence, the rostral but not the cZI was reported as visible (Kerl et al., 2013), which we demonstrate is actually the fasciculus lenticularis (Figure 2). As a result, protocols for stereotactic targeting of the cZI have relied on the relative visibility of the surrounding RN and STN, from which the location of the stereotactic target within the PSA could be indirectly inferred. Overall, our results confirm that the ZI is not a strong generator of T2w contrast and led us to explore other potential generators of MR contrast.

In the present study, we found that T1 rather than T2w relaxation properties of the ZI better delineated the substructures in the region. T1 relaxation times increase in a field-dependent manner, as does the dispersion between brain tissue types (Rooney et al., 2007), which have the effect of improving contrast between tissue types at 7T. This advantage has been exploited to parcellate thalamic nuclei (Tourdias et al., 2014) and investigate cortical laminae (Trampel et al., 2017). Surgical planning and *in vivo* histology have been considered important potential applications of the MP2RAGE sequence (Marques et al., 2010; Marques and Gruetter, 2013). In fact, using this method, we demonstrate that the ZI can also be visualized along its entire rostrocaudal axis (Figure 1). Furthermore, we found sufficient difference in T1-related tissue parameters to permit separation of the cZI from surrounding white matter tracts, including the fct of the PSA and the fields of Forel (ft and fl) from the rZI (Figure 2). Rostrally, these contrast differences permitted more detailed characterization of the relationship between the fl and rZI, dividing the rZI into dorsal and ventral components described in experimental animals (Mitrofanis, 2005; Watson et al., 2014) and one human brain atlas (Mai et al., 2015). Although the increase in T1 tissue values with field strength has been perceived as a disadvantage due to increased scan time, our results indicate that sufficient resolution and contrast can be attained within a clinically reasonable timeframe.

Since the boundaries of the ZI have not previously been well-defined in three dimensions, consensus segmentations were performed using group averaging to further boost the SNR when delineating these structures. Our interpretation of the boundaries of the ZI using *in vivo* sequences was based on detailed comparison with annotations of the ZI from classical and modern histological sections (Hawrylycz et al., 2012; Mai et al., 2015; Morel, 2007; Schaltenbrand and Wahren, 1977). The majority of the segmented structures in the ZI region could be reliably segmented in both the template space and for individual scans (Dice > 0.70 being considered reliable) although segmentation of the rZI, fl, and ft in individual subjects was less reliable (Table 3). Dice scores are generally lower for smaller structures, as small random errors in the boundary have a larger relative weight when volumes are smaller. For the rZI, the complex morphology of this region and its relationship with the white matter tracts of the fields of Forel is likely another contributing factor. Whether to include the newly identified (and previously unlabeled) region between H2 and the STN also likely increased uncertainty of segmentation of the rZI. We opted via consensus to include this in the definition of the rZI although this will have to be investigated in future studies integrating histology and other methods. For the fl and ft the decreased reliability likely relates to the small size of these structures (50 mm^3^ and 80 mm^3^) as well as the challenge of identifying the lateral limits of segmentation given that they are white matter structures projecting to other nuclei. To compute estimates across the study population, the template segmentations were propagated back to the individual datasets using the transformations computed during the template creation process. This template creation approach allowed for the pooling of data from multiple participants (N=32) into a single reference space allowing us to better account for intersubject variability. Compared to histological evaluations, our approach enables high-resolution imaging without the drawbacks of histological processing, which include tissue deformations, processing artifacts, and other technical issues (Morel, 2007; Nowacki et al., 2018b).

Our analysis demonstrates that there is sufficient signal and contrast within the PSA region to allow separation of the cZI from the fct (see Table 4). We discovered that commonly used T2w indirect anatomical target and optimal stimulation locations appeared at the boundary of the cZI with the PSA (Figure 4 and Supplementary Figure S4). These findings are in line with other work suggesting that a proportion of benefit is derived from stimulation of wayward white matter tracts in the fct (raprl) (Blomstedt et al., 2018; Mohadjer et al., 1990; Mundinger, 1965; Spiegel et al., 1964; Velasco et al., 1972), and also concordant with recent studies employing diffusion tensor imaging (DTI) (Dallapiazza et al., 2018; Dembek et al., 2019; Fenoy and Schiess, 2017; Fiechter et al., 2017; Sammartino et al., 2016; Velasco et al., 2018). Compared to DTI-based measures, ultra-high field T1 mapping has higher SNR, is less prone to image distortions, require generally less scan time, less post-processing, and is acquired at inherently higher resolution (0.7 mm isotropic compared to 2-3 mm). We have determined that the dimensions of the fct within the PSA is ∼4-5 mm along its longest axis, representing 1-3 voxels if relying on DTI alone compared to 5-7 voxels using our protocol. Direct visualization presents the possibility of submillimetric to millimetric level refinement of the therapeutic target and stimulation parameters, particularly if newer current steering devices are implanted.

Our findings add to the growing body of knowledge that the optimal DBS target within the PSA is at the anterior boundary of the cZI abutting or directly within the fasciculus cerebellothalamicus (Fiechter et al., 2017; Herrington et al., 2016). This suggests that direct targeting of the white matter, in other words connection-based targeting, may be central to efficacy, which has increasingly been acknowledged for essential tremor (Akram et al., 2018; Al-Fatly et al., 2019) and other disorders (Horn et al., 2017). Our approach using T1 mapping for visualizing local WM tracts might be considered divergent from recent approaches using diffusion-based imaging. With respect to human *in vivo* studies, DTI studies have mostly focused on connections between larger cortical and subcortical structures since achieving high resolution (submillimetric) images in clinically feasible timeframes for DTI remains a challenge. There is also increasing acknowledgement that connectivity-based methods are prone to producing false-positive tracts (Maier-Hein et al., 2017). An additional advantage of using T1 mapping, is that the images can simultaneously be used as a baseline structural scan and furthermore used to identify the target, eliminating the need for an image fusion step, which can introduce error. Ultimately, the approach taken here, particularly with increasingly higher resolution imaging, should be considered complementary to diffusion-based endeavors, enabling accurate localization of smaller tracts and structures using a multi-contrast approach. For example, anatomical segmentations of local white matter tracts at the template and individual participant levels could be used to optimize tractographic and connectomic approaches, as seed regions to boost sensitivity to smaller tracts.

Some discrepancy in T1 map values was noted when comparing values reported between sites (Figure 3b) and studies (Forstmann et al., 2014; Keuken et al., 2017). In particular, our values tended to be ∼100-200 ms shorter within the STN and SN. Ideally, quantitative maps should be independent of imaging sites and scanner vendors, and indicative of underlying tissue parameters. However, several factors may account for discrepancies between studies employing comparable quantitative imaging approaches (Stikov et al., 2015). For quantitative T1 mapping, the inversion recovery (IR) method is traditionally considered the gold standard (Drain, 1949; Hahn, 1949). However, several limitations, including long scan times associated with the acquisition of many images, reduce its utility for practical purposes. Therefore, more time efficient methods like the DESPOT (Deoni et al., 2005) and MP2RAGE (Marques et al., 2010) sequences have gained significantly in popularity in the last decade, with the latter being commonly acquired for higher field strength T1 mapping. In contrast to the traditional IR approach, the MP2RAGE approach requires the acquisition of only two images at different inversion times, which due to the interleaved nature of the sequence are inherently co-registered. This limits the effect of subject motion on the precision to map T1 and delineation of subcortical regions. In addition, whereas more conventional anatomical sequences applied in the clinic are influenced by M0 (i.e., proton density), T2*, B1- (i.e. radiofrequency [RF] receive) and B1+ (i.e., transmit) fields, the MP2RAGE approach removes these effects by only varying the inversion time and flip angles between each inversion image. However, we have recently shown that the slight variations in MP2RAGE setup between the original and replication datasets can introduce strong variability of cortical T1 across the brain, with observed differences leading up to 260 ms between datasets (Haast et al., 2020). These differences are most presumably related to differences in their sensitivity to B1+ inhomogeneities as post-hoc B1+ correction (Eggenschwiler et al., 2012; Marques and Gruetter, 2013) lowered inter-dataset offsets to under 100 ms for both cortical (Haast et al., 2020), as well as subcortical T1 (this paper). Moreover, inter-scanner variability in hardware – our use of parallel versus single RF transmission for tissue excitation – may amplify this B1+ sensitivity. In addition, the differences in acquisition parameters (Supplementary Table S5) can introduce additional sequence-dependent measurement variability due to assumptions about mono-exponential T1 relaxation in the MP2RAGE implementation (Rioux et al., 2016). Although differences in T1 are observed between the original and replication datasets, a striking correspondence is visible in terms of the relative T1 values between assessed regions proving the value of T1 mapping to identify these regions in a time efficient manner. Finally, how the findings in this study hold in the presence of pathology or atrophy remains an unanswered question and will be the subject of future work.

## Conclusions

In the present study, we demonstrate that direct *in vivo* visualization of the structures of the human ZI region is possible, a region originally described as an “immensely confusing area about which nothing can be said.” We successfully derived estimates of the size, shape, location, and tissue characteristics of substructures in the peri-zonal region non-invasively at high (submillimetric) resolution. Our findings confirm observations, only previously possible through histological evaluation, that the ZI is not simply a space between structures but contains distinct morphological entities that should be considered separately. Our findings pave the way for increasingly detailed *in vivo* study and provide a structural foundation for precise functional and neuromodulatory investigation bringing increasing certainty to this uncertain area.

## Supporting information

Video: template space and labels (sagittal orientation)

Video: template space and labels (coronal orientation)

Video: template space and labels (axial orientation)

## Acknowledgements

JL is funded through the Western University Clinical Investigator Program accredited by the Royal College of Physicians and Surgeons of Canada and a CIHR Frederick Banting and Charles Best Canada Graduate Doctoral Award Scholarship. KU was supported by a grant from the Institute for Basic Science, Suwon, Republic of Korea (IBS-R015-D1). The work is supported by postdoctoral fellowships from BrainsCAN to RH and YX, and CIHR to YX. Support from CIHR Foundation grant FDN 201409 is also acknowledged. We would like to thank Catherine Currie for her assistance with recruiting participants for this study.

