## Supplementary Materials for "Direct visualization and characterization of the human zona incerta and surrounding structures"

### Supplementary Sections

#### Supplementary Section S1. Nomenclature

**Supplementary Table S1.** Nomenclature of the zona incerta region used in this study

| **Study nomenclature*** | **Abbreviation** | **Synonyms** |
|---|---|---|
| zona incerta | ZI | |
| caudal zona incerta | cZI | |
| rostral zona incerta | rZI | |
| fasciculus thalamicus | ft | thalamic fasciculus, H1 field of Forel |
| fasciculus lenticularis | fl | lenticular fasciculus, H2 field of Forel |
| field H | fh | H field of Forel<br>mergence of fasciculus lenticularis and ansa lenticularis (Gallay et al., 2008) |
| medial lemniscus | ml | lemniscus medialis |
| fasciculus cerebellothalamicus | fct | radiatio prelemniscus / prelemniscal radiations (raprl) / brachium conjunctivum<br>dentatorubrothalamic tract (drtt) |
| subthalamic nucleus | STN | corpus subthalamicum, corpus Luysi |
| red nucleus | RN | ruber, nucleus ruber tegmenti |
| posterior subthalamic area** | PSA | |

* nomenclature generally follows the naming scheme used by Morel (Morel, 2007) and also Schaltenbrand (Schaltenbrand and Wahren, 1977)

** term used in the stereotactic neurosurgery literature to refer to a functional target for treating tremor

### Supplementary Section S2. Data curation and template validation

**General Results:**

The Anatomical Fiducials (AFIDs) framework was used for annotating key point features in each of the 32 participants (Lau et al., 2019). Anatomical fiducial locational errors (AFLE), in other words, placement errors were computed. Mean AFLE after quality control was 0.91 +/- 0.69 mm. Visual inspection of AFIDs revealed good spatial correspondence as well as evidence of good anatomical detail in subcortical structures of the final template (Figure 1). T2w-to-T1w image registration failed in 2 out of 32 participants (6.25%) during image preprocessing. With manual intervention, we were able to correct the registration for these two subjects. Ultimately, the iterative deformable template creation process resulted in a convergence to a mean AFID registration error of 1.27 +/- 1.02 mm after 10 iterations (Figure 1b). Mean AFRE decreased globally across all AFID points with registration complexity improving from 29.62 +/- 11.58 mm at baseline to 3.47 +/- 1.62 mm after linear registration and to 2.80 +/- 0.90 mm after deformable registration (one iteration). Further increases were noted with successive iterations of template generation to 2.06 +/- 0.92 mm after 4 iterations to 1.27 +/- 1.02 mm after 10 iterations. Mean AFRE improved to a limit of 2.82 +/- 1.47 mm with linear registration alone with little improvement beyond 6 iterations, while improvements were noted up to 10 iterations with deformable registration (Figure 1b and Supplementary Figure 1).

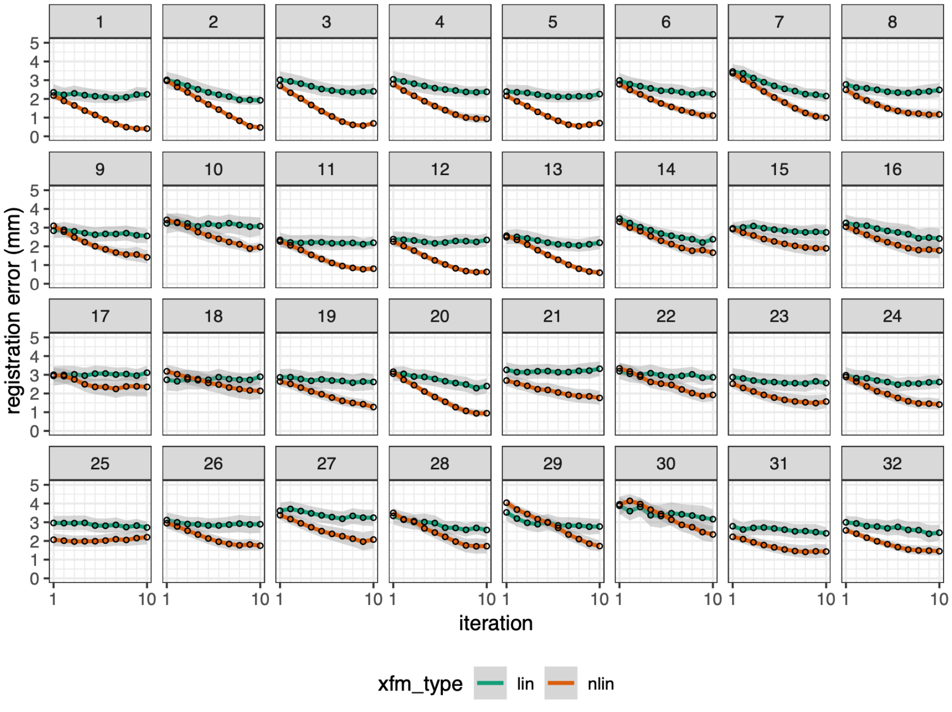


**Supplementary Figure S2. Registration error evaluation for validated anatomical fiducials.** All the features improved with multiple iterations and the addition of nonlinear registration methods. Several features also improved to the submillimetric range, noting particularly AFID01 (anterior commissure) and AFID02 (posterior commissure). See Lau et al. for more details on the corresponding categories for each AFID (Lau et al., 2019).

### Supplementary Section S3. Manual segmentation results for RN and STN.

**Supplementary Table S3.** Intra- and inter-rater reliability for segmenting known regions.

| | | Intra-Rater | | Inter-Rater | |
|---|---|---|---|---|---|
| **roi** | **side** | **jaccard** | **kappa** | **jaccard** | **kappa** |
| **RN** | left | 0.95±0.03 | 0.98±0.02 | 0.92±0.04 | 0.96±0.02 |
| **RN** | right | 0.95±0.03 | 0.98±0.01 | 0.91±0.03 | 0.95±0.02 |
| **STN** | left | 0.90±0.10 | 0.94±0.06 | 0.81±0.07 | 0.90±0.04 |
| **STN** | right | 0.89±0.10 | 0.94±0.05 | 0.79±0.04 | 0.88±0.02 |

#### Supplementary Section S4. Posterior subthalamic area plots

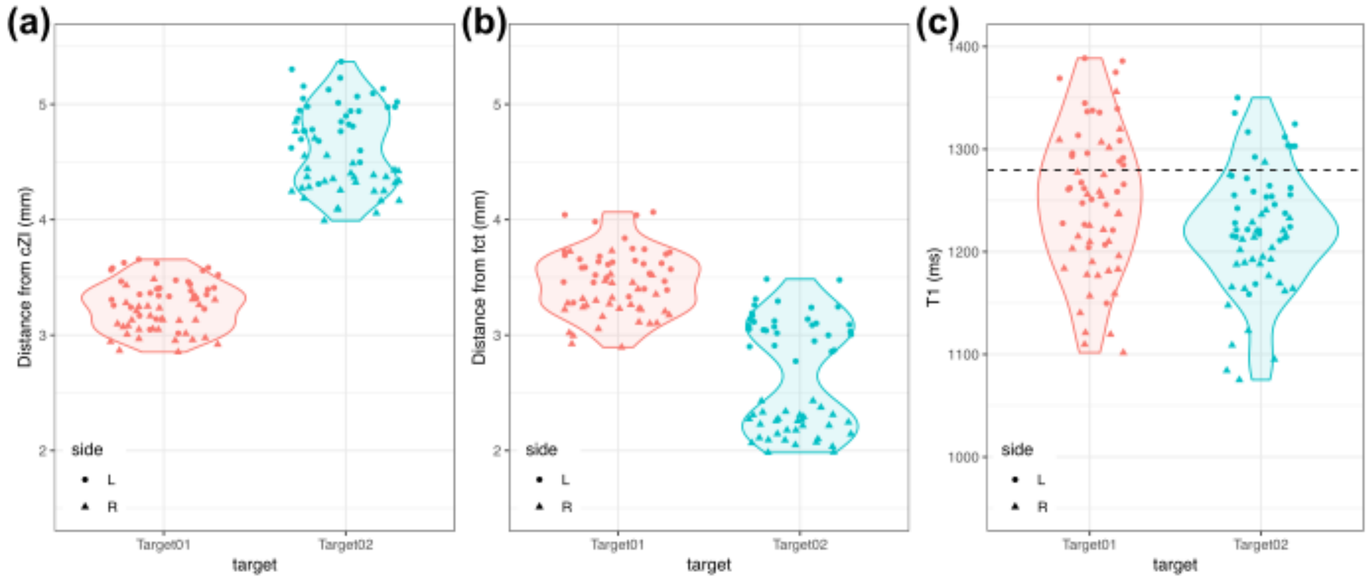


**Supplementary Figure S4. The relationship between the indirect targets and distance from nearby structures of the zona incerta region and T1 values.** (a+b) Demonstration that Target01 (Blomstedt et al., 2010) is about equidistant from cZI and fct whereas Target02 (Nowacki et al., 2018a) is further away from cZI but closer to fct. (c) T1 values in both targets are slightly lower than the mean value in the cZI (dashed line; Figure 4 and Table 3) with Target 02 slightly lower. Together these provide objective evidence that existing indirect targets are at boundary regions between these two structures.

### Supplementary Section S5. Comparison of MP2RAGE acquisition data

**Supplementary Table S4.** Comparison of different protocols for MP2RAGE sequence at 7T.

| Sequence | | TE (ms) | TR (ms) | TI (ms) | Flip Angle (°) | Matrix Size | PAT* | Averages | Resolution ($mm^3$) | Acquisition Time (s) |
|---|---|---|---|---|---|---|---|---|---|---|
| Current Study | 3D | 2.73 | 6000 | 800/2700 | 4/5 | 342x342x224 | 3 | 1 | 0.7x0.7x0.7 | 10:14 |
| Replication Data* | 3D | 2.47 | 5000 | 900/2750 | 5/3 | 320x320z240 | 3 | 1 | 0.7x0.7x0.7 | 08:02 |
| Forstmann 2014 | 3D | 2.45 | 5000 | 900/2750 | 5/3 | 320x320x240 | 2 | 1 | 0.7x0.7x0.7 | 10:57 |

* from Maastricht University Brain Imaging Centre

### Supplementary Section S6. Study template with labels.

**Video S6a.** Video fly through of the study template with label overlays on the left side and no overlay on the right (axial orientation). Red = red nucleus, green = subthalamic nucleus, blue = zona incerta, yellow = fasciculus cerebellothalamicus, magenta = fields of Forel.

**Video S6b.** Video fly through of the study template with label overlays on the left side and no overlay on the right (coronal orientation). Red = red nucleus, green = subthalamic nucleus, blue = zona incerta, yellow = fasciculus cerebellothalamicus, magenta = fields of Forel.

**Video S6c.** Video fly through of the study template with label overlays on the left side and no overlay on the right (sagittal orientation). Red = red nucleus, green = subthalamic nucleus, blue = zona incerta, yellow = fasciculus cerebellothalamicus, magenta = fields of Forel.
